# Structure-activity of outer membrane proteins in native bacterial membrane vesicles by solid-state NMR

**DOI:** 10.64898/2026.07.15.738770

**Authors:** Tata Gopinath, Kyungsoo Shin, Swapna Bera, Alyssa Kraft, Karthika Samimuthu, Ashish K. Gadicherla, James E Kent, Nicholas A Wood, Francesca M. Marassi

## Abstract

The outer membranes (OMs) of bacterial pathogens are potent virulence factors and serve as the first line of defense against host immunity. Their striking bilayer asymmetry is essential for function but poses an exceptional challenge for reconstitution *in vitro*, limiting structure-activity analysis to artificial non-native platforms that can interfere with structure and function. Here, we describe bacterial OM vesicles (OMVs), natively secreted from the cellular OM during bacterial cell growth and development, as an effective vehicle for structure-activity analysis *in situ* based on solid-state nuclear magnetic resonance (NMR). We show that *E. coli* OMVs may be engineered to express a range of isotopically labeled target OM proteins, and isolated for solid-state NMR magic angle spinning (MAS) experiments. High resolution NMR spectra are obtained for three bacterial virulence factors: the adhesion invasion locus (Ail) and plasminogen activator protease (Pla) from *Yersinia pestis*, and the major porin (OmpF) from *E. coli*. The spectra reflect the native protein structures, report on the specific OMV membrane environment, and may be used to map protein interactions with their human host ligands, specifically the multifunctional glycoprotein Vitronectin (Vn) which binds Ail as part of its serum protection activity. Notably, OMVs support protein functionality, enabling structure and activity to be correlated *in situ*. OMVs expressing plasmid-encoded Ail recruit human Vn and confer serum protection to wild-type *E. coli* cells, while OMVs expressing plasmid-encoded Pla support the proteolytic activity of Pla. Taken together, the data establish OMVs as a robust new platform for structure-activity analysis of OM proteins *in situ*, offer new insights about the complexity of the bacterial OM, and reveal additional functional aspects of OMVs as key ancillary units of bacterial infection.

## Introduction

The outer membranes (OMs) of diderm bacteria mediate first-line processes that are critical for promoting bacterial cell survival and dissemination in the human host, including resistance to complement-mediated lysis, adhesion to human cells, biofilm formation, suppression of blood clotting, and more. Bacterial OMs, including their lipid and protein components, are potent virulence factors and attractive targets for therapeutic drug development, but their complex architecture and striking asymmetry ^1-4^ pose an exceptional challenge for *in vitro* reconstitution, and this has limited atomic-level structural characterization to formulated platforms – detergent micelles, bicelles, nanodiscs and liposomes – that do not capture the properties of the native OM and can alter structure and activity ^5^.

In the bacterial OM, phospholipids and lipopolysaccharide (LPS) molecules are each segregated to the outer and inner leaflets, and have co-evolved with OM proteins for specialized inside-outside membrane functionalities that provide both a potent permeability barrier and the first line of defense against immunogenic attack from the human host. They have mutually stabilizing effects important for maintaining integrity of the bacterial cell envelope ^6^, and increasing evidence points to the OM as a coordinated proteolipid assembly where LPS molecules are shared by multiple proteins ^7,8^. This complexity elevates the importance of generating structure-activity data *in situ*.

Here we introduce biogenic bacterial OM vesicles (OMVs) as an excellent platform for structure-activity studies based on solid-state nuclear magnetic resonance (NMR). OMVs are natively secreted spherical nanoscale structures that originate from the cellular OM. Extracellular vesiculation is a regulated OM remodeling mechanism distinct from either stress-induced explosive cell lysis or cell death ^9-12^, and OMVs perform many fundamental biological functions including the delivery of virulence factors, environmental adaptation, host survival and colonization, and biofilm formation. Important for structure-activity analysis, OMVs can be harvested directly from bacterial cell cultures, and preserve the native asymmetric architecture of the bacterial OM while bypassing the need for purification, detergent treatment or reconstitution. Moreover, their closed vesicular architecture supports the correct transmembrane display of OM proteins, enabling external ligands to be added for structure-activity analysis.

Working with three proteins with key roles in microbial virulence, and sizes ranging from 8 to 10 and 16 stranded β-barrels – the adhesion invasion locus Ail and plasminogen activator protease Pla from *Yersinia pestis*, and the major porin OmpF from *E. coli* (**Fig. 1A**) – we show that OMVs can be engineered and isolated directly for NMR-based structure-activity analysis. The resulting NMR spectra are highly resolved. They reflect the complex environment of the bacterial OM and reveal NMR chemical shift perturbations that correlate with the specific OMV membrane environment and map interactions with human host ligands, specifically the interaction of Ail with the multifunctional glycoprotein Vitronectin (Vn). Importantly, OMVs also support the functional activities of serum protection activity of Ail and the proteolytic activity of Pla, enabling structure and activity to be correlated *in situ*. The work introduces OMVs as a new platform for NMR structure-activity studies of bacterial OM proteins in native membranes.

**Figure 1.**
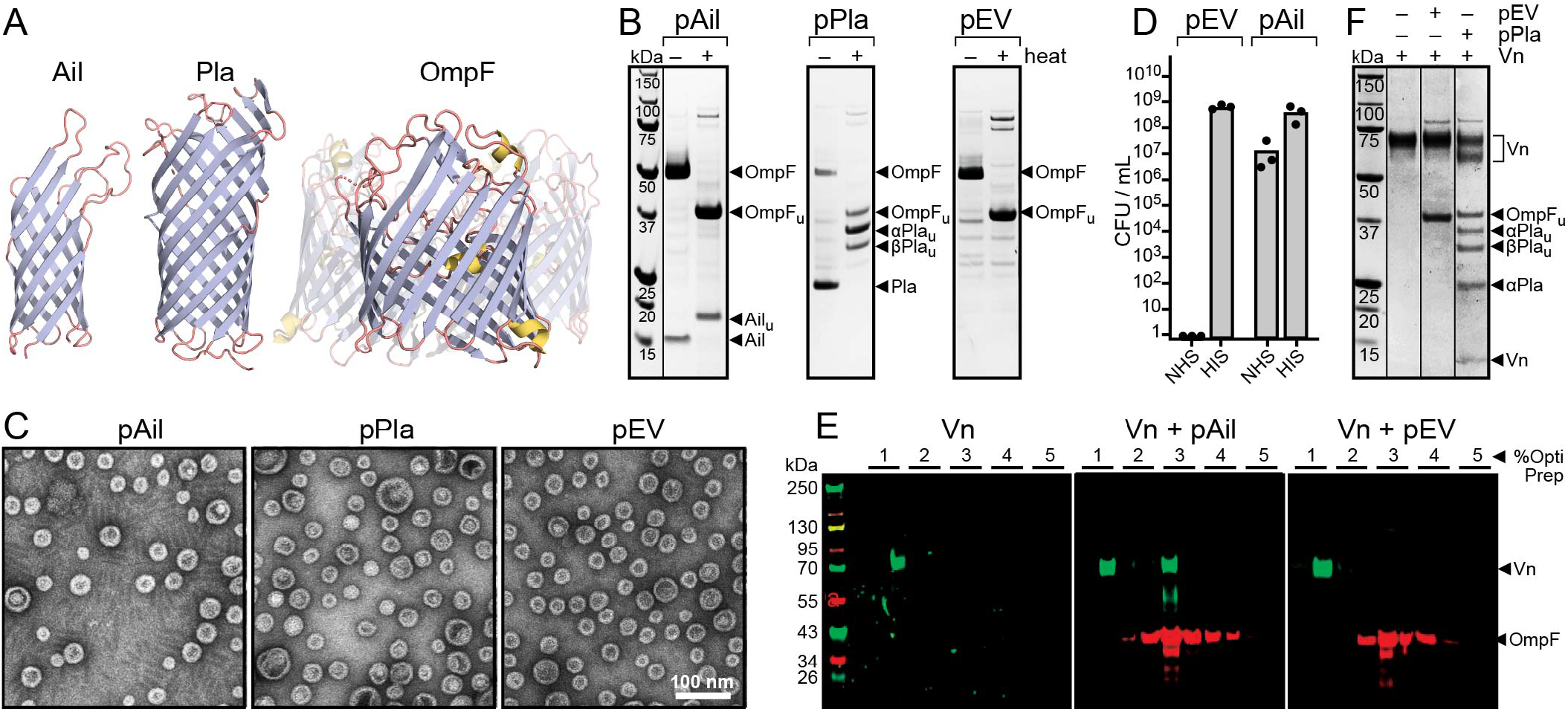
Ail, Pla and OmpF localize to *E. coli* OMVs. **(A)** Structures of Ail (PDB 2N2L; ^26^), Pla (PDB 2X55; ^24^), and OmpF (PDB 2ZFG; ^27^). **(B)** SDS-PAGE of OMVs isolated from pAil, pPla, and pEV (empty plasmid) *E. coli* cell cultures. Proteins were visualized with Coomassie stain. Arrows mark bands from heat-unfolded (u) or folded proteins. **(C)** Representative negative stain EM of pAil, pPla and pEV OMVs. **(D)** Serum protection activity of pAil OMVs. Each bar represents the average number of colony forming units (CFUs) per mL of cell culture, measure over three replicate experiments. *E. coli* cells were incubation with NHS or HIS supplemented with either pEV or pAil OMVs, then serially diluted and plated on LB-agar supplemented with ampicillin (100 µg/mL) and chloramphenicol (35 µg/mL). After incubating at 37°C overnight, the bacterial colonies were enumerated to estimate the number of CFUs per mL. **(E)** Western blot of density gradient fractions obtained for Vn alone, Vn added to pAil OMVs, or Vn added to pEV OMVs. **(F)** Proteolytic activity of pPla OMVs towards human Vn.

## Results and Discussion

### Ail, Pla and OmpF localize to bacterial OMVs

*Y. pestis* Ail was one of the first OM proteins to be directed to OMVs by heterologous expression in *E. coli* ^13^. Ail enhances the virulence of the class A pathogen *Y. pestis* by promoting cell adhesion and invasion, biofilm formation and bacterial resistance to the immune defenses present in human serum ^14-18^. While Ail is an abundant component of the cellular OM proteome, *Y. pestis* also releases OMVs enriched in Ail, Pla, and other virulence factors, as a mechanism that is thought to influence the outcome of infection by facilitating pathogen dissemination ^19^.

Previously, we showed that Ail can be targeted to the OM of engineered *E. coli* for direct solid-state NMR studies in the native bacterial cell envelope ^20,21^. The NMR spectra reflect the 8-stranded β-barrel structure and reveal sites that interact with LPS, other cell envelope components, and human serum components. Moreover, since *E. coli* support the serum resistance, auto-aggregation, and Vn binding activities of Ail, they serve as a useful model system for structure-activity analysis *in situ*.

Here, we relied on the same engineered *E. coli* approach to induce the expression of both isotope-labeled Ail and Pla in the bacterial OM but took biogenic OMVs isolated directly from the media of the bacterial cell cultures for NMR analysis ^20,22,23^. Induction of plasmid-encoded Ail (pAil) or Pla (pPla) results in a significant amount of either protein localizing to OMVs, as evidenced by SDS-PAGE (**Fig. 1B**). The Ail β-barrel migrates as a ∼15 kDa band that shifts to ∼18 kDa once unfolded by heat, as observed previously in whole cells ^20^. Similarly, the 10-stranded β-barrel of Pla ^24^ migrates as a ∼24 kDa band that separates into two ∼30 and ∼34 kDa bands with heating, as expected for the auto-processed enzyme ^25^. No evidence of protein misfolding was detected in either pAil or pPla OMVs.

In both cases, an intense ∼50 kDa band that shifts to ∼37 kDa upon heating is observed. This species is also abundant in OMVs isolated from cells transformed with empty plasmid vector (pEV) but otherwise treated as pAil and pPla cells, and was previously assigned to the 16-stranded OmpF porin by mass spectrometry and immunoblotting ^28^. The relative intensities of the Ail, Pla and OmpF bands in OMVs compared to the cell fraction, indicate that Pla is highly directed to OMVs compared to Ail (**Fig. S1**). All three OMV types (pAil, pPla, pEV) are spherical and homogeneously sized (∼35 nm), with the inner and outer leaflets of the single membrane visibly resolved as electron-dense regions by electron microscopy (**Fig. 1C**).

### Bacterial OMVs support the activities of *Y. pestis* Ail and Pla

In a previous study ^29^, OMVs isolated from *Salmonella enterica* were shown to promote bacterial cell survival in serum by interfering with complement activation. This activity was linked to PagC, an Ail homolog, and major promoter of OM vesiculation, that localizes to OMVs ^29-31^ when upregulated by the two-component PhoP-PhoQ system under conditions of Ca^2+^ and Mg^2+^ ion starvation, low pH, or cationic antimicrobial peptides ^32,33^. Ail is also known to promote the resistance of *Y. pestis* to complement-dependent killing by human serum ^34^, and plasmid-encoded Ail expression has been shown to render *E. coli* DH5α and Lemo-21(DE3) cells immune to serum mediated killing ^20,35^, but the role of OMVs in this regard has not been explored. Here, we find that *E. coli* pAil OMVs can confer serum resistance when added to *E. coli* cell cultures (**Fig. 1D**). *E. coli* Lemo-21(DE3) cells survive the bactericidal activity of normal human serum (NHS) when supplemented with pAil but not pEV OMVs, compared to control experiments where cells are incubated with heat inactivated serum (HIS).

Ail has also been shown to bind the human serum protein vitronectin (Vn) ^35^, an important fluid-phase regulator of both the complement and coagulation pathways. Vn regulates innate immunity through its interaction with membrane attack complex components, and also interacts with PAI-1 (plasminogen activator inhibitor 1) stabilizing its function as a key brake in the conversion of plasminogen to plasmin during fibrinolysis. To test the ability of pAil OMVs to bind Vn, we performed density gradient sedimentation experiments with glycosylated Vn expressed and purified from HEK293 mammalian cells (**Fig. S2**). The data show that Vn co-sediments with pAil OMVs but not pEV OMVs, leading us to conclude that pAil OMVs are capable of recruiting Vn (**Fig. 1E**).

Pla is a plasmid-encoded Asp protease expressed at the *Y. pestis* cell surface ^24,36,37^. Pla processes plasminogen to generate active plasmin protease which then degrades fibrin, and Pla also degrades Vn ^35^ and the Vn/PAI-1 complex ^38^, thereby activating the dissolution of blood clots and extracellular matrix networks and promoting systemic pathogen dissemination and invasion ^39^. We find that pPla OMVs, but not pEV OMVs, efficiently degrade purified Vn (**Fig. 1F**), yielding a proteolytic product of similar in size to the 65 kDa form of Vn, as reported previously for whole *Y. pestis* cells ^35^.

Notably, *Y. pestis* has been shown to release OMVs that contain both Ail and catalytically active Pla and can bind various human host factors ^19^, and the dissemination of Ail and Pla via secreted OMVs is thought to potentiate *Y. pestis* infection and virulence. The finding that pAil OMVs confer serum resistance to wild-type bacterial cells adds to the evidence that OMV production can play an important role in neutralizing the innate immune defenses of the host and supports a role for OMVs in promoting the local and systemic spread of bacteria in human infections.

### Engineered bacterial OMVs yield high-resolution solid-state NMR spectra

To obtain protein-specific ^15^N and ^13^C isotopic labeling for NMR studies, the T7/DE3-mediated expression of pAil and pPla was induced with IPTG, and bacterial cells were grown in ^15^N/^13^C labeled minimal media coupled with suppression of chromosomal gene transcription by the antibiotic rifampicin. For OmpF, isotopic labeling could be obtained without either plasmid-engineered expression or rifampin treatment, in line with its abundance as the dominant chromosomally-encoded OM protein in pEV OMVs. For all three proteins, the biogenic OMVs were harvested directly from the growth media and centrifuged into 3.2 mm or 1.3 mm rotors for solid-state NMR experiments at 14 kHz or 60 kHz magic angle spinning (MAS) frequencies.

The resulting two-dimensional ^1^H/^15^N and ^15^N/^13^C and NMR spectra are highly resolved (**Fig. 2; Fig. S3**). In the spectra of Ail (156 residues; **Fig. 2A, B**), several assignments can be tentatively transferred from the assigned spectra (BMRB: 30284) in liposomes and nanodiscs ^40,41^, indicating that the overall structure of the 8-stranded β-barrel is supported in all these environments. Moreover, direct comparison with the spectra acquired for the cell envelope fractions of the parent bacterial cell cultures reveals extensive overlap, as expected for these very similar OM environments (**Fig. 2 red**). We also observe several new signals in the OMV spectra compared to cell envelope, micelles, nanodiscs and liposomes ^6,26,40,41^, most notable in the resolved spectral region associated with Gly (**Fig. 3A-C, Fig. S4**). The majority of invisible signals in the spectra from micelles, nanodiscs and liposomes ^6,26,40,41^ are associated with the extracellular loops, where conformational disorder in non-native platforms leads to broadened and low-intensity NMR lines, as well as incomplete observation of electron density in X-ray diffraction. While resonance assignments of the OMV spectra will be needed, the appearance of additional signals suggests that the OMV platform provides an environment more supportive of loop conformational order and conducive to NMR analysis.

**Figure 2.**
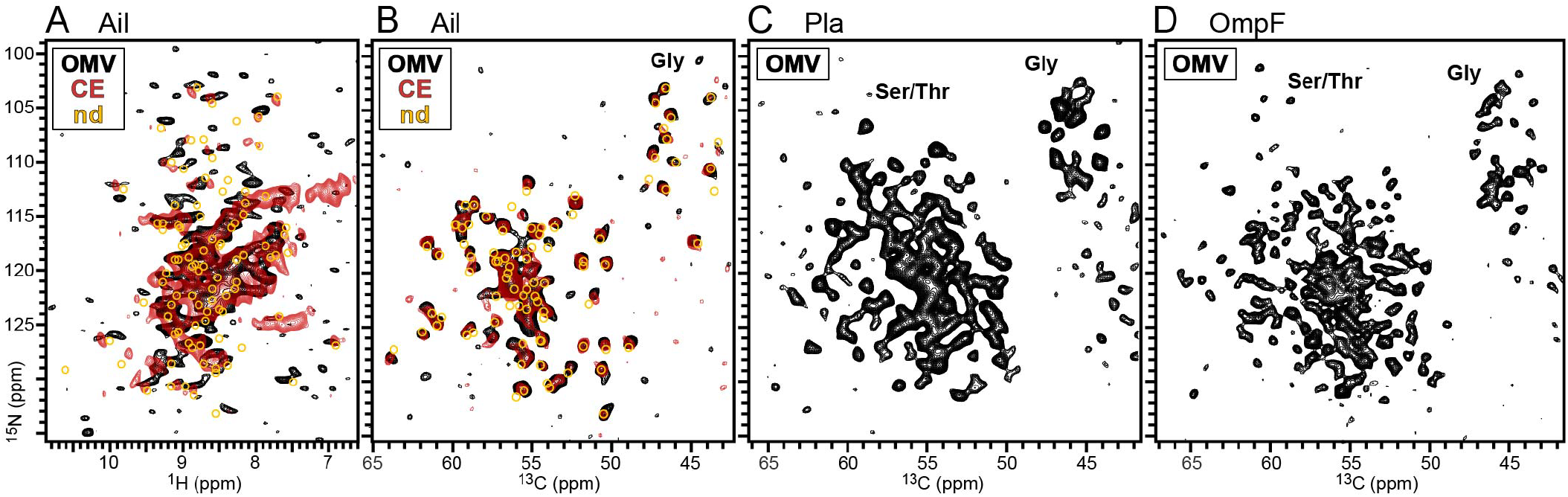
Two-dimensional solid-state NMR spectra of native *E. coli* OMVs. Spectra were acquired for OMVs (black) or the cell envelope fraction (CE; red). Previously assigned spectra ^40^ were obtained for purified Ail reconstituted in nanodiscs by solution NMR (nd; yellow circles). **(A)** ^1^H/^15^N CP-HSQC spectra of Ail. The cell envelope spectrum was recorded previously at 900 MHz with 57 kHz MAS ^20^. **(B-D)** ^15^N/^13^C NCA spectra of Ail, Pla, and OmpF. The cell envelope spectrum (red) was recorded in this study.

**Figure 3.**
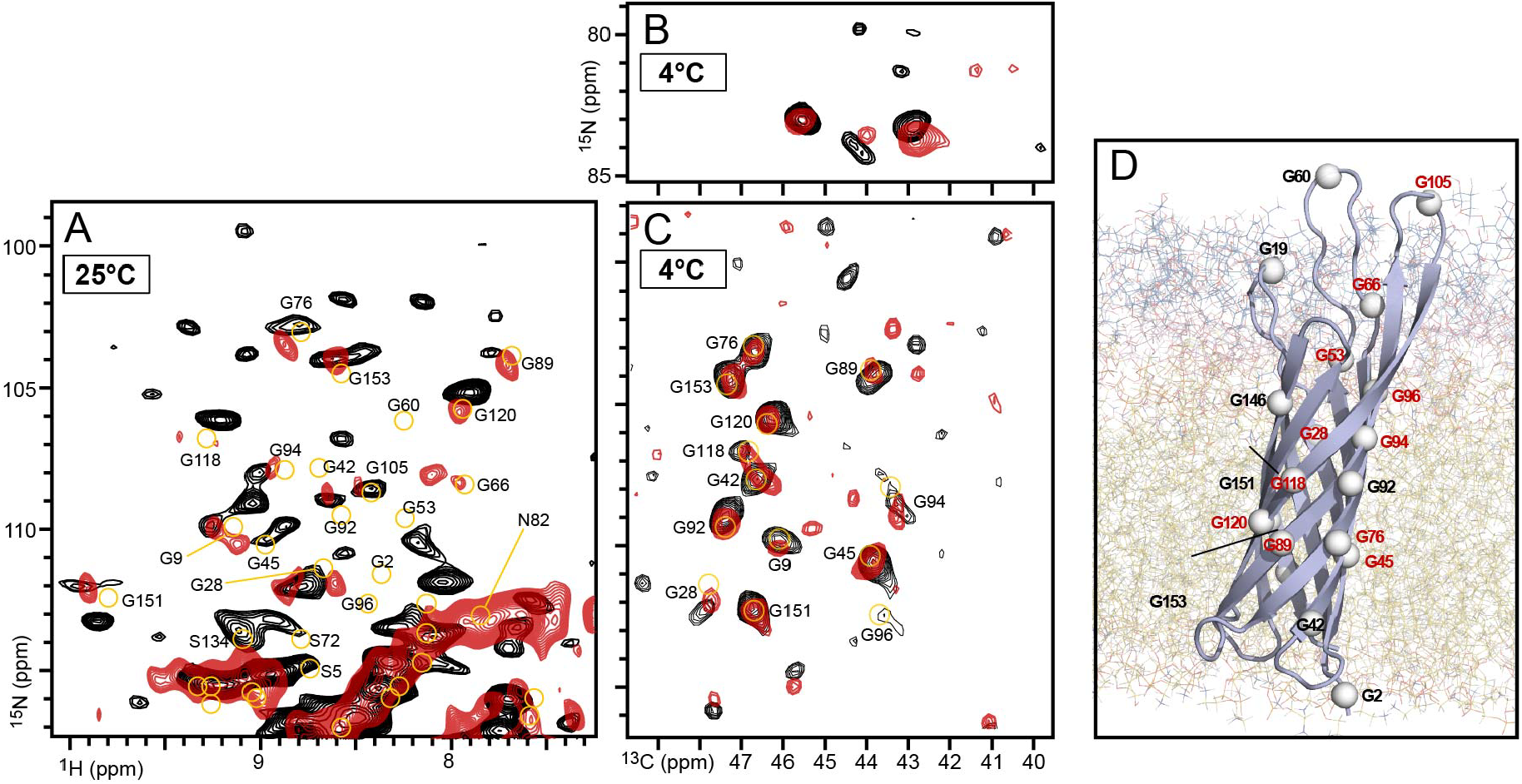
NMR spectra of Ail reveal differences between the OMV and cell envelope environments. **(A-C)** Expanded regions of the two-dimensional ^1^H/^15^N CP-HSQC and ^15^N/^13^C NCA spectra of Ail in biogenic OMVs (black) or bacterial cell envelope (red). Assignments represent the solution NMR spectra of Ail in nanodiscs (yellow circles) ^40^. The cell envelope ^1^H/^15^N spectrum was recorded previously at 900 MHz with 57 kHz MAS ^20^ (A). The cell envelope ^15^N/^13^C spectra (red) were recorded in this study (B, C). **(D)** Structural model of Ail embedded in the *Y. pestis* OM taken from previous MD simulation ^6^. Spheres denote the 20 Gly N atoms. Sites with different ^1^H/^13^C/^15^N chemical shifts in OMVs versus cell envelope are labeled red. The OM is shown as lines with LPS outer core (blue), LPS inner core (pink), and LPS and PL acyl chains (yellow).

In the ^15^N/^13^C spectra of Pla (252 residues) and OmpF (340 residues), the region associated with Gly (^13^C: 42-48 ppm; ^15^N: 100-120 ppm) resolves approximately 24 of the 26 Gly signals expected for Pla, and 35 of the 48 Gly signals of OmpF. Notably, the spectrum of OmpF also resolves several signals in the ^15^N region associated with the Ser and Thr backbone (^15^N: 100-105 ppm). These are significantly shifted upfield relative to the average values, as expected for the few Ser and Thr residues located in the highly shielded electrostatic environment of the OmpF conducting pore. These residues play important roles in regulating porin selectivity ^42,43^ and the ability to detect and resolve their NMR signals offers new opportunities for probing the OmpF porin activity *in situ*.

### NMR reveals differences between OMV and cell envelope OM environments

Since OMVs have the same asymmetric OM architecture of their parental bacterial cells, we expected the OMV and cell envelope spectra of Ail to be identical when acquired at the same temperature and MAS frequency, but some residue-specific chemical shift differences are observed (**Fig. 3A-C; Fig. S4**). Those from resolved spectral regions that may be assigned with some confidence map across the length of the Ail β-barrel, including sites embedded within the OM (**Fig. 3D**).

The largest chemical shift differences are observed for ^1^H and ^15^N, as expected from the high sensitivity of these nuclei to the local environment. The ^15^N/^13^C and ^1^H/^15^N spectra were each acquired at sample temperatures of 4°C and 25°C, as our spectrometer is not capable of lowering the sample temperature below 25°C at a MAS frequency of 60 kHz, and some of the differences between the OMV and cell envelope spectra are observed at 25°C and not 4°C. For example, the G76, G89, G120 ^15^N signals are shifted upfield by ∼0.5ppm in OMVs compared to cell envelope (**Fig. 3A**) but do not appear to change significantly at 4°C (**Fig. 3B**).

Inspection of the Arg guanidinium spectral region also reveals chemical shift differences and the appearance of new peaks in OMVs compared to cell envelope (**Fig. 3B**). Signals from the Arg sidechain NE-CD atoms are resolved from the backbone by means of Hadamard editing ^44^, and for the eight Arg residues in the Ail sequence, eight NE-CD signals are observed in the OMV spectrum compared to six in cell envelope. Three Arg at the intracellular membrane surface (R34, R80, R155) are buried in the barrel interior, and the other five located at the extracellular surface (R14, R27, R51, R52, R110) have been shown to interact with LPS polar groups ^21^. Observation of their NMR signals suggests that they possess a degree of conformational order in both OMVs and cell envelope, while the chemical shift differences point to a different Ail-LPS interaction in the two environments.

Taken together, the NMR data show that Ail is sensitive to distinct OMV and cell envelope environments, suggesting that they may differ in significant ways despite their common origin. What might be the cause of this difference? The OMV membranes can differ in composition from their parent cells ^9-12^, and various physical properties of the OM, including lateral pressure, conformational dynamics and protein-protein clustering, may be expected to differ significantly in the extreme membrane curvature structures of native OMVs (∼30-35 nm diameter) relative to the relatively planar structures of the cell envelope. For example, OM proteins and LPS molecules have been shown to cluster in distinct islands with restricted mobility, rather than freely diffusing across the OM surface, ^45-48^, and both diffusivity and lateral heterogeneity may be expected to differ in the high-curvature membranes of OMVs compared to the parent cells.

### Interaction of Ail with Vn

Mapping the interaction of OM proteins with their human host ligands is a long-standing goal for advancing structure-guided therapeutic development. Since NMR chemical shifts are highly susceptible to the local environment, they can report on even very weak intermolecular interactions that cause minimal structural perturbation, and thus offer an effective approach for structure-activity analysis. The OMV platform is ideally suited for these experiments because its natively closed structure retains the asymmetry of the bacterial OM with protein extracellular regions displayed on the OMV surface and intracellular regions occluded inside the lumen. This architecture is distinct from nanodiscs, which are completely open with both membrane leaflets equally exposed to the same aqueous environment, and it is also different from cell envelope fragments, where the membranes artificially reseal in ways that can occlude protein extracellular regions.

To map the Ail-Vn interaction, we co-sedimented purified Vn with ^15^N/^13^C pAil OMVs into the MAS rotor for NMR experiments. Comparison of the ^1^H/^15^N spectra of pAil OMVs obtained with or without purified Vn reveal several ^1^H/^15^N chemical shift perturbations (**Fig. 4, Fig. S5**). Mapping these changes to specific residues is challenging without direct experimental assignments of the ^1^H/^15^N spectra, but to the extent that some assignments may be transferred from the nanodisc spectra, they appear to map across the Ail β-barrel (**Fig. 4C**).

**Figure 4.**
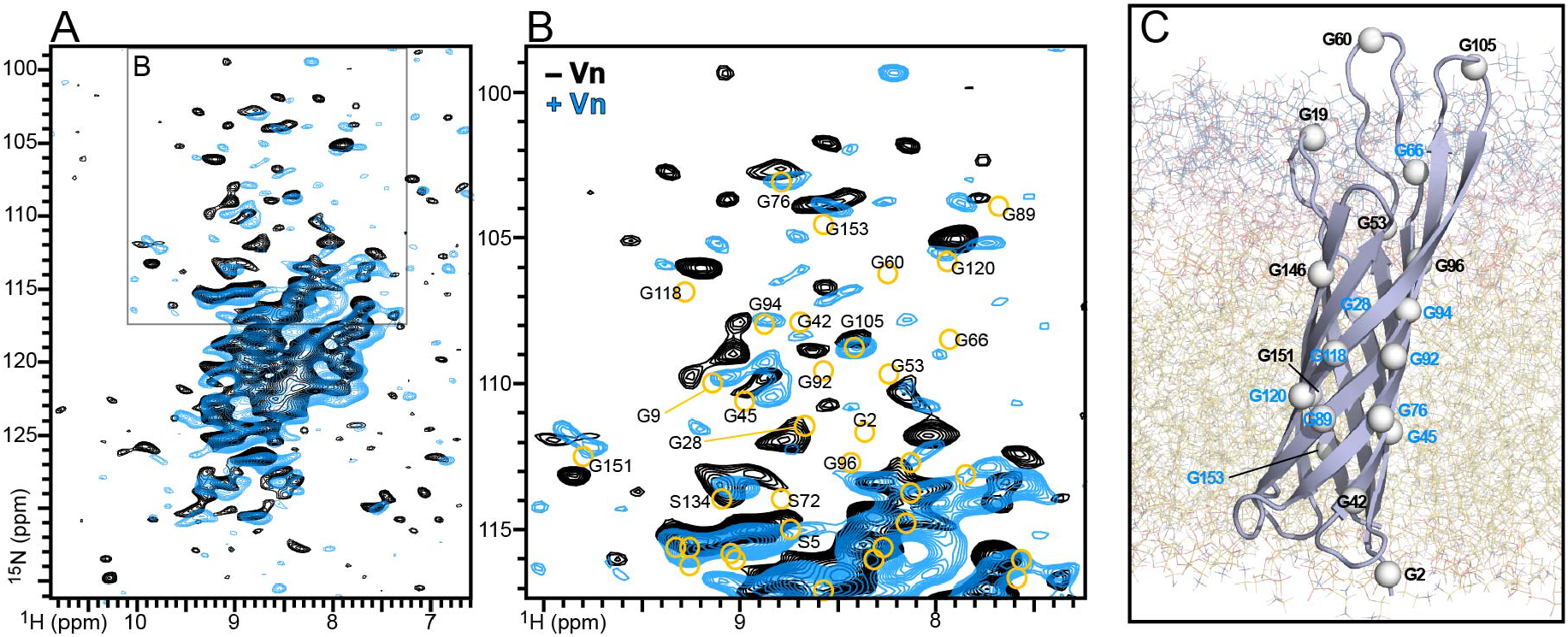
NMR ^1^H/^15^N spectra of Ail reveal differences between the OMV and cell envelope OM environments. **(A, B)** Two-dimensional ^1^H/^15^N CP-HSQC spectra of Ail in biogenic OMVs sedimented without (black) or with purified Vn (blue). Previously assigned spectra solution NMR, shown for comparison, were obtained for purified Ail reconstituted in nanodiscs (yellow circles) ^40^. Inset box denotes expanded spectral area (B). **(C)** Structural model of Ail embedded in the *Y. pestis* OM taken from previous MD simulation ^6^. Spheres denote amide N atoms with tentatively assigned chemical shifts that are perturbed by addition of Vn. The OM is shown as lines with LPS outer core (blue), LPS inner core (pink), and LPS and PL acyl chains (yellow).

Ail is essential for recruiting Vn to the bacterial OM ^35^, and residues in its extracellular loops (F54, F68, S102, and F104) have been proposed to form a binding site critical for binding Vn ^49^. Signals from G53 and G94, near the proposed Vn binding site, appear to be perturbed by Vn, but ^1^H/^15^N perturbations are also observed for the intramembrane region of the Ail β-barrel. There are two possible explanations for this result. The association of Vn with the extracellular loops of Ail could cause conformational changes that transmit allosterically to membrane-embedded sites of the Ail β-barrel, or the NMR perturbations could reflect perturbations of the OM environment caused by the Ail-Vn interaction. The two mechanisms are not mutually exclusive and may both contribute to a global alteration of the OM induced by Vn association, which itself could be related to the serum protection activity associated with Vn binding.

### OMVs represent an effective new platform for NMR structure-activity studies *in situ*

OMVs play many fundamental biological functions and have important roles as vaccine, drug delivery and nanotechnology vehicles ^9-12^. OMVs purified from wild-type bacteria have been used to examine the roles of virulence factors in the host, while OMVs from engineered bacterial strains represent an important tool for studying protein functions *in situ* ^13^. Recently, we showed that OMVs can be used for NMR structural studies to gain mechanistic insights about PagC-driven OMV biogenesis ^28^. In this study, we extend their application to a broad range of OM proteins for NMR-based structure-activity analysis. We show that bacterial OMVs support native protein activities and yield high-resolution NMR spectra that enable protein-OM and protein-protein interactions to be mapped with atomic resolution. The finding that Ail OMVs protect bacterial cells from the bactericidal activity of human serum is particularly interesting and strengthens the evidence for an important role of OMVs in immune evasion.

OMVs offer a number of advantages compared to other cellular platforms. They can be directly harvested from the culture media, bypassing the need for membrane fractionation and treatment with detergents, and they preserve the asymmetric architecture of the bacterial OM. Their natively sealed spherical architecture is ideal for analyzing ligand binding at the outer surface of the OM. While engineered overexpression can lead to the accumulation of misfolded OM protein in the periplasmic space that co-purifies with the cell envelope fraction, OMVs appear to incorporate exclusively folded proteins, an important advantage for the detection of homogeneous single-line NMR spectra Moreover, the NMR spectra from OMVs are free of background signals from peptidoglycan and other cell envelope components. Taken together, the present results offer new insights about the complexity of the bacterial OM, reveal additional functional aspects of OMVs as ancillary functional units in bacterial infections, and introduce OMVs as a new platform for NMR structure-activity studies *in situ*.

## Methods

### Bacterial strains, culture conditions, and key reagents

All *E. coli* were commercially available Lemo21(DE3) (New England BioLabs, MA, USA). The pET22b(+)-*ail* (pAil) and pET22b(+)-*pla* (pPla) plasmids were designed for IPTG-inducible expression of Ail or Pla, each lacking their endogenous signal peptide and in frame with the PelB signal peptide ^50^, and acquired from Genscript (Piscataway, NJ, USA). Bacteria were transformed with empty pET22b(+) plasmid (pEV) vector for control experiments. Bacteria were cultured in M9 media supplemented with 0.2% glucose, 100 µg/mL ampicillin and 35 µg/mL chloramphenicol when appropriate. Protein expression was monitored by sodium dodecyl sulfate (SDS) polyacrylamide electrophoresis (PAGE).

### Isotope labeling

Transformed bacteria were grown overnight at 37°C in unlabeled M9 media. Overnight cultures were used to inoculate fresh unlabeled M9 to achieve OD_600_=0.1, and cultures were incubated with shaking at 37°C. At OD_600_=0.6, expression of the DE3 RNA polymerase was induced by adding 0.4 mM IPTG. Following a 10-minute incubation, the cells were pelleted (5,000xg, 4°C, 20 min), and resuspended in M9 supplemented with 1 g/L of ^15^N ammonium sulfate and 0.2% ^13^C glucose (Cambridge Isotope Laboratories, MA, USA). For selective labeling, IPTG and rifampin were added to 0.4 mM and 100 µg/mL, respectively. Expression was allowed to proceed for 16 hours at 26° C prior to harvesting of samples. For OmpF isotope labeling, pEV cells were cultured as the pAil and pPla cells without adding either rifampin or IPTG.

### OMV isolation and characterization

OMVs were harvested by diafiltration and ultracentrifugation as described ^28^. Briefly, all cultures were centrifuged repeatedly to remove bacterial cells, the supernatant was filtered through a 0.45 µm membrane, and then concentrated using an Amicon stirred cell fitted with a 300 kDa PES membrane disc filter (Millipore Sigma, St. Louis, MO, USA). OMVs were harvested by centrifugation (150,000 g, 4°C), washed by resuspension in buffer (0.9 mM CaCl_2_ and 0.5 mM MgCl_2_), and finally harvested for analysis by centrifugation (150,000 g, 4°C). OMVs were quantified by light scattering and UV absorbance, as described ^51,52^.

### Bacterial cell envelop preparation

Cell envelope fractions were isolated from pAil cells as described ^20,21^. Briefly, the cells were isolated by centrifugation, then lysed by three passes through a French Press, and the cell envelope fraction harvested by centrifugation after washing with buffer (20 mM NaH_2_PO_4_/Na_2_HPO_4_, pH 6.5).

### Negative stain EM

Purified OMVs (3 *µ*L; 1 mg/mL) were adsorbed onto glow-discharged 400 mesh Formvar carbon copper grids (Electron Microscopy Sciences, PA, USA) for 15 sec, and then wicked away with filter paper. The grids were then treated with 2% uranyl-acetate for 30 sec, then blotted to remove the stain, and allowed to dry for at least one hour prior imaging at 100,000x magnification on a Jeol JEM-1400 Transmission Electron Microscope (JEOL USA, MA) at 60 kV.

### Expression and purification of Vitronectin

The pcDNA3.1 expression vector encoding the sequence of human Vn (residues 1-478) was designed to include a 5’ Kozak consensus sequence and a C-terminal Gly_6_-His_8_ tag, and acquired from by Genscript (Piscataway, NJ, USA). The vector (pVn-GH) was transfected into Expi293 cells (Gibco), and the cells cultured in Expi293 expression medium with shaking (37°C, 5% CO_2_, 80% humidity, and 130 RPM) following standard procedures ^53^. Protein expression was allowed to continue for 3 days before harvesting the culture supernatant by centrifugation (1,000 g, 10 minutes, 4 °C), and adjusting the solution to the final resuspension buffer concentration (20 mM Tris-HCl, pH 7.8, 300 mM NaCl). Vn was isolated from the buffered supernatant by Ni-affinity chromatography with Ni-Excel resin (Cytiva). Vn was eluted with imidazole (300 mM), and further purified by size-exclusion chromatography using a Superdex S-200 10/300 GL increase column (Cytiva) in resuspension buffer. Purified vitronectin was stored at 4°C. . Protein expression and purification were monitored by SDS PAGE and western blot with αVn (209-258; R12-2413, Assay Biotech) and αHis (Qiagen) antibodies.

### Serum Protection experiments

Serum-susceptible pEV *E. coli* was used to inoculate M9 minimal medium. After overnight culture, the cells were transferred to fresh M9 (starting OD_600_ = 0.1) and then incubated with shaking at 37°C, to reach OD_600_ = 0.5. Cells were harvested by centrifugation and washed with sterile PBS (three times) before resuspension in half the original volume of PBS to achieve OD_600_ =1.0. For assays, the washed cells (100 µL) were transferred to a deep-well 96 well culture plate, mixed with purified OMVs (300 µg in 50 µL PBS) by gentle pipetting, and finally supplemented with 50 µL of either normal human serum (NHS) or heat-inactivated serum (HIS). The assay plates were covered with a gas-permeable seal, incubated with shaking (37°C, 150 rotations per minute) for 30 minutes, and then placed on ice, before each sample was serially diluted, plated on fresh LB agar plates, and incubated overnight at 37°C. Colonies were enumerated the next day, and colony counts were used to back-calculate colony forming units (CFUs) per mL.

### Pla activity assay

pPla OMVs or pEV OMVs were suspended at 0.05 mg/mL in buffer (20 mM Tris-HCl pH 7.8, 300 mM NaCl) and incubated with purified Vn at 4°C overnight. Vn cleavage by Pla was assayed by SDS PAGE.

### Solid-state NMR spectroscopy

Purified OMVs or cell envelope fractions were centrifuged into 3.2 mm or 1.3 mm zirconium rotors. Solid state NMR experiments were performed on a 700 MHz NeO Bruker spectrometer, using a 3.2 mm BlackFox ^1^H/^13^C/^15^N probe optimized for ^13^C detection, or a 1.3 mm ^1^H/^13^C/^15^N probe optimized for ^1^H detection. The effective sample temperature under MAS conditions was estimated by measuring the water ^1^H resonance frequency. NMR spectra were processed with NMRpipe and analyzed using Poky (Sparky) software ^54,55^. NMR experimental details are provided in Table S1.

The ^15^N/^13^C/hNCA spectra were acquired at a sample temperature of 4°C, with MAS rates of 14 kHz, or 12.5 kHz, and a recycle delay of 2 s. The 90° pulse lengths were set to 3 µs (^1^H), 6 µs (^13^C) and 6 µs (^15^N). ^1^H-^15^N cross polarization was obtained with ^15^N radiofrequency amplitudes of 30 kHz or 35 kHz. The Hartmann-Hahn matching condition was set by ramping the ^1^H radiofrequency amplitude to maximize signal intensity. Double ^1^H-^15^N-^13^C cross polarization transfer was obtained with radiofrequency spinlocks of 23 kHz (^13^C) and 35 kHz (^15^N), and 100 kHz continuous wave decoupling (^1^H). Heteronuclear decoupling was achieved using the spinal-64 pulse sequence with 95 kHz radiofrequency amplitude. The ^1^H-detected ^1^H/^15^N CP-HSQC spectra were acquired at a sample temperature of 25°C, with a MAS rate of 60 kHz, and a recycle delay of 2 s, as described previously ^20,41^

## Data Availability

Source data for all figures presented in the paper and Supplementary Information are available.

PDB 2N2L: https://www.rcsb.org/structure/2N2L

PDB 2X55: https://www.rcsb.org/structure/2X55

PDB 2ZFG: https://www.rcsb.org/structure/2ZFG

BMRB 30284: https://bmrb.io/data_library/summary/index.php?bmrbId=30284

## Acknowledgments

This work was supported by grants from the National Institutes of Health (GM118186 and AI188770). It utilized NMR instrumentation supported by a grant from the NIH (OD028716), and transmission EM instrumentation supported by the MCW-Oxford Instruments Center for Advanced Microscopy and Electron Microscopy Core (RRID: SCR_026315). We thank Marassi lab members for discussion.

## Author contributions

TG, KS, JEK, NAW and FMM were involved in research design, data analysis and discussion. TG, KS, JEK, SB, AK, KS, AKG, JEK and NAW performed experiments. FMM wrote the manuscript with input and approval from all authors.

