## Supplementary data for "Structure-activity of outer membrane proteins in native bacterial membrane vesicles by solid-state NMR"

**Table 1. NMR Samples and solid-state NMR experimental parameters.** Data were acquired on a 700 MHz Bruker spectrometer under magic angle spinning (MAS), for outer membrane vesicles (OMV) or cell envelope (CE) samples.

| Sample | Experiment and MAS Frequency | Cross Polarization Contact Time | Number of Transients | t1 Acquisition Time | t2 Acquisition Time | Experiment time |
|---|---|---|---|---|---|---|
| **pEV OMV pH 7** | NCA 12.5 kHz | HN-CP 2 ms<br>NCA-CP 5 ms | 2048 | 12.8 ms | 15 ms | 97 h 20 m |
| **pPla OMV pH 7** | NCA 14 kHz | HN-CP 1ms<br>NCA-CP 4 ms | 2048 | 15.7 ms | 15 ms | 114 h |
| **pAil CE pH 7** | NCA 14 kHz | HN-CP 1.5 ms<br>NCA-CP 4 ms | 2560 | 14.7 ms | 15 ms | 137 h |
| **pAil OMV pH 7** | NCA 14 kHz | HN-CP 1.5 ms<br>NCA-CP 4 ms | 2560 | 14.7 ms | 15 ms | 137 h |
| | hNH 60 kHz | HN-CP 1.2 ms<br>NH-CP 0.5 ms | 4096 | 14.8 ms | 8 ms | 101 h, 20 m |
| **pAil OMV + Vn pH 7** | hNH 60 kHz | HN-CP 1.2 ms<br>NH-CP 0.5 ms | 2048 | 14.8 ms | 8 ms | 110 h, 47 m |

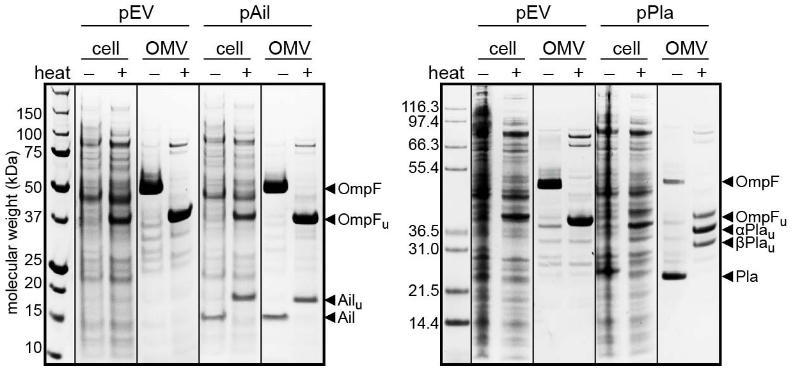


**Figure S1. Localization of Ail, Pla and OmpF to OMVs and cellular OM.** SDS-PAGE of whole cells and OMVs isolated from pAil, pPla, and pEV (empty plasmid) *E. coli* cell cultures. Proteins were visualized with Coomassie stain. Arrows mark bands from folded proteins and heat-unfolded (u) proteins.

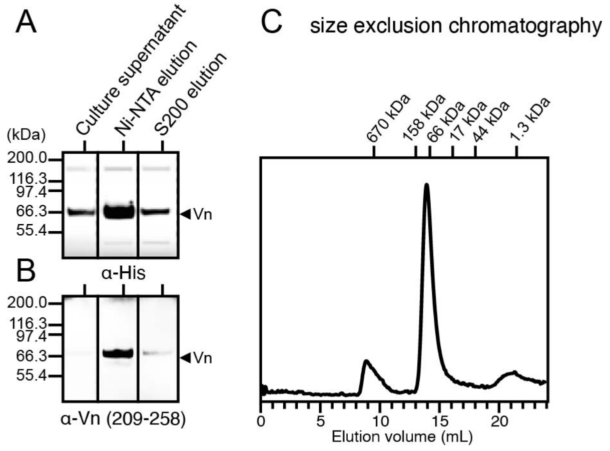


**Figure S2. Expression and purification of native human Vn from HEK293 cells. (A, B)** Western blot analysis of Vn fractions with anti-Vn(209-258) and anti-His antibodies. **(C)** Size exclusion chromatography on a Superdex S-200 10/300 GL increase column (Cytiva) yields homogeneously pure protein.

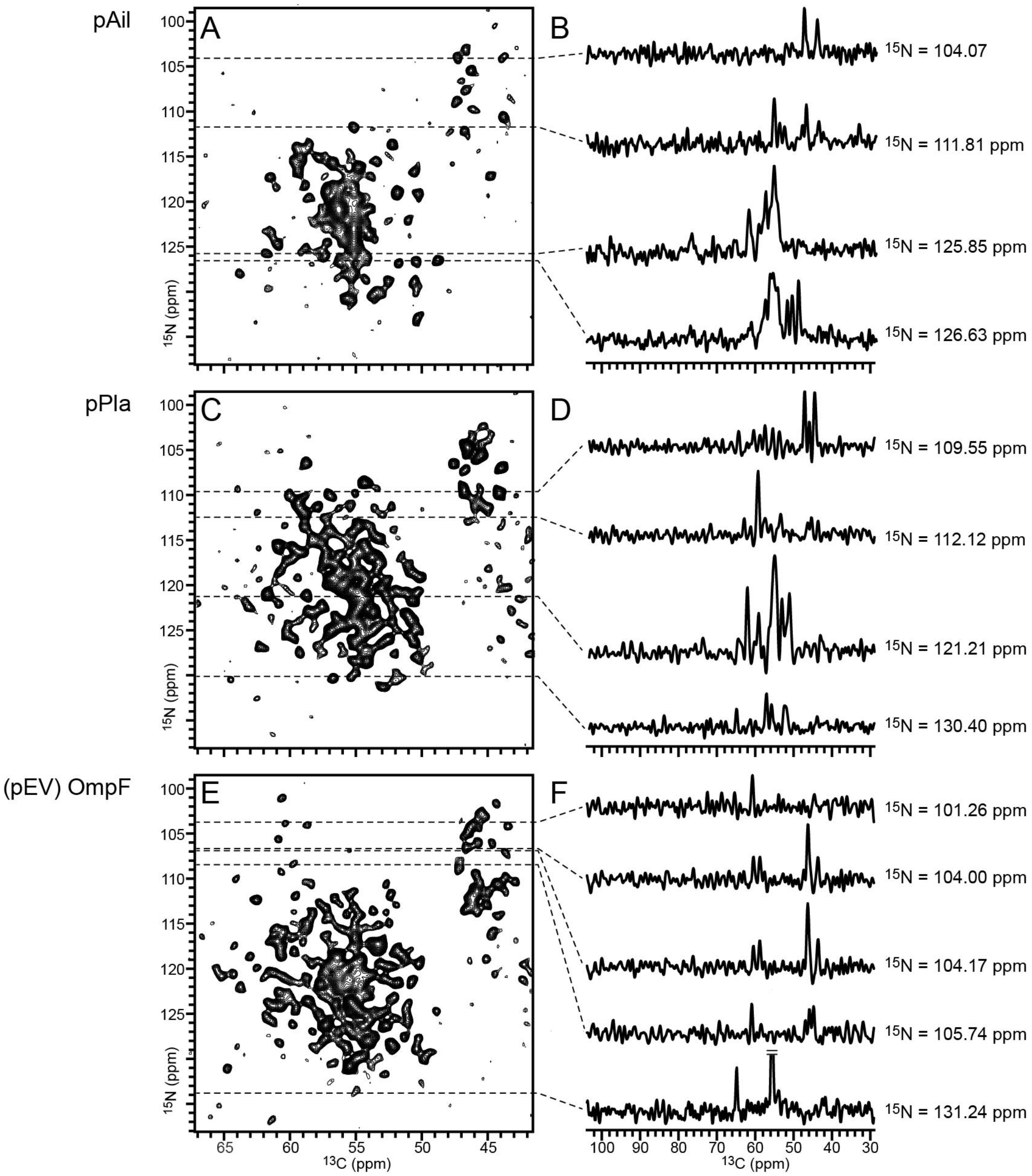


**Figure S3. Two-dimensional $^{15}$N/$^{13}$C solid-state NMR spectra of Ail, Pla and OmpF in bacterial OMVs. (A, C, E)** Two-dimensional $^{15}$N/$^{13}$C NCA spectra. **(B, D, F)** One-dimensional spectra taken from the 2D spectra at $^{15}$N chemical shifts marked by the dashed lines.

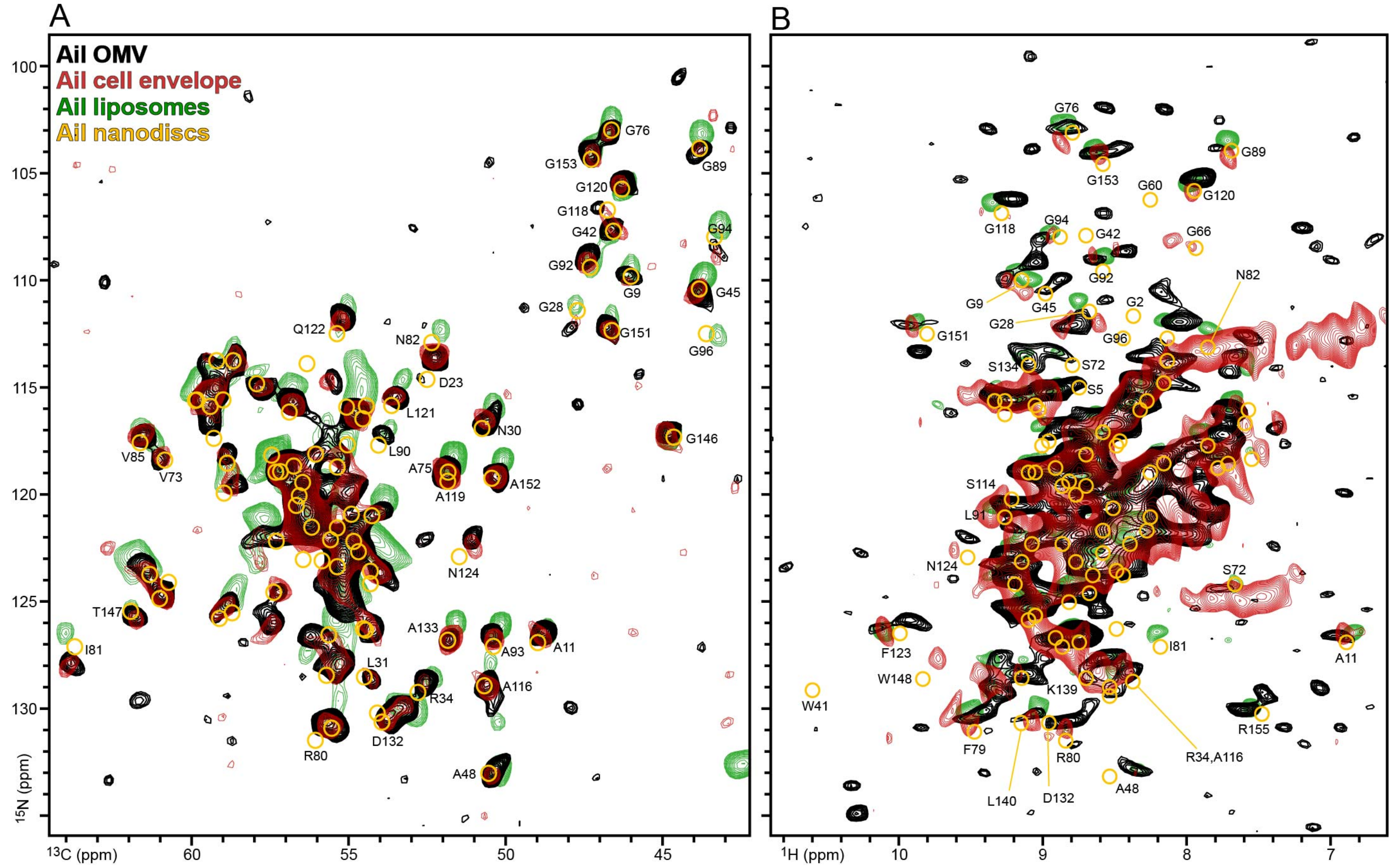


**Figure S4. Two-dimensional solid-state NMR spectra of Ail in the native bacterial membrane.** Spectra were acquired for pAil OMVs (black) or pAil cell envelope (red). Previously assigned spectra were obtained for purified Ail reconstituted in liposomes by solid-state NMR (green) [41] or in nanodiscs by solution NMR (yellow circles) [40]. **(A)** Solid-state NMR $^{15}$N/$^{13}$C NCA spectra. The cell envelope $^{15}$N/$^{13}$C spectrum (red) was recorded in this study. **(B)** Solid-state NMR $^{1}$H/$^{15}$N CP-HSQC spectra. The cell envelope spectrum was recorded previously at 900 MHz with 57 kHz MAS [20].

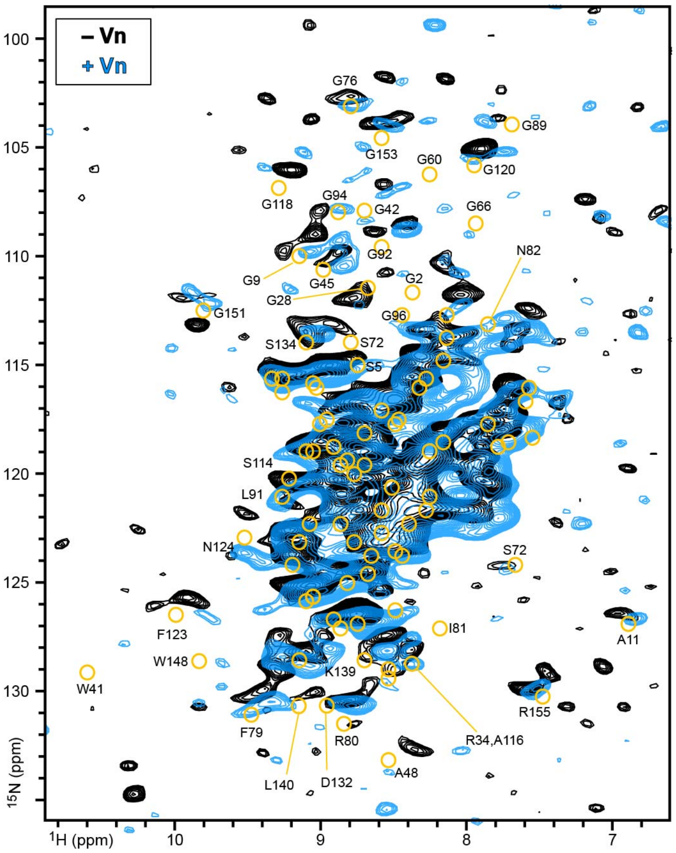


**Figure S5. Two-dimensional solid-state NMR $^{1}$H/$^{15}$N CP-HSQC spectra of pAil OMVs.** Spectra were acquired for pAil OMVs (black) or pAil OMVs co-sedimented with purified Vn (blue). Previously assigned solution NMR spectra were obtained for purified Ail reconstituted in nanodiscs (yellow circles) [40].
